# Genomic prediction using individual-level data and summary statistics from multiple populations

**DOI:** 10.1101/314609

**Authors:** Jeremie Vandenplas, Mario P.L. Calus, Gregor Gorjanc

**Affiliations:** Wageningen University & Research, Animal Breeding and Genomics, 6700 AH Wageningen, The Netherlands; The Roslin Institute and Royal (Dick) School of Veterinary Studies, University of Edinburgh, Easter Bush Research Centre, Midlothian EH25 9RG, UK

**Keywords:** meta-analysis, quantitative trait, statistical method

## Abstract

This study presents a method for genomic prediction that uses individual-level data and summary statistics from multiple populations. Genome-wide markers are nowadays widely used to predict complex traits, and genomic prediction using multi-population data is an appealing approach to achieve higher prediction accuracies. However, sharing of individual-level data across populations is not always possible. We present a method that enables integration of summary statistics from separate analyses with the available individual-level data. The data can either consist of individuals with single or multiple (weighted) phenotype records per individual. We developed a method based on a hypothetical joint analysis model and absorption of population specific information. We show that population specific information is fully captured by estimated allele substitution effects and the accuracy of those estimates, i.e. the summary statistics. The method gives identical result as the joint analysis of all individual-level data when complete summary statistics are available. We provide a series of easy-to-use approximations that can be used when complete summary statistics are not available or impractical to share. Simulations show that approximations enables integration of different sources of information across a wide range of settings yielding accurate predictions. The method can be readily extended to multiple-traits. In summary, the developed method enables integration of genome-wide data in the individual-level or summary statistics form from multiple populations to obtain more accurate estimates of allele substitution effects and genomic predictions.

## INTRODUCTION

Genome-wide markers are nowadays widely used to predict complex traits. This prediction is based on a linear model that partitions for each individual the observed complex phenotype value into systematic effects, comprising at least a population mean, an individual genetic value and an environmental deviation (Fisher, 1918). With genome-wide markers, individual genetic values can be computed from allele substitution effects estimated from individual-level phenotype and genotype data (Meuwissen et al., 2001). Subsequently, genetic values can be also computed for individuals of interest that are genotyped, but not phenotyped. This process is commonly called genomic prediction. In animal and plant breeding, genetic values are used to identify genetically superior individuals and use them as parents of the next generation to improve complex traits like milk yield (Meuwissen et al., 2001; VanRaden, 2008) or grain yield (Schulthess et al., 2016) In human genetics, genetic values can be used to predict individual genetic risk for complex diseases to inform preventive and personalized medicine (Campos et al., 2010; Wray et al., 2013; Pasaniuc and Price, 2017).

Accuracy of estimated allele substitution effects and of resulting genetic values for complex traits are foremost a function of the amount of available data (Daetwyler et al., 2008). To maximize the prediction accuracy, use of all available data is recommended (Henderson, 1984; Wray et al., 2013; Vilhjálmsson et al., 2015). In some small populations, collecting large amounts of data is not possible, and a joint analysis across multiple populations is needed to achieve high accuracy (Hozé et al., 2014; Wientjes et al., 2016). However, such joint analysis is often impossible, because of logistic or privacy considerations (Powell and Norman, 1998; Maier et al., 2018). Therefore, several methods were proposed to enable analysis of data from multiple populations when individual-level data is not available (Pasaniuc and Price, 2017; Liu and Goddard, 2018; Maier et al., 2018). These methods approximate a joint analysis by first obtaining summary statistics from separate analyses of individual-level data for each population and then combine these summary statistics to estimate genetic values. In human genetics, summary statistics usually consist of publically available allele substitution effects, i.e., genome-wide associations, together with their standard errors, estimated independently for each marker (Yang et al., 2012; Vilhjálmsson et al., 2015; Maier et al., 2018). In livestock, summary statistics more likely consist of allele substitution effects estimated jointly for all markers, together with prediction error (co)variances (Liu and Goddard, 2018). While these methods may increase prediction accuracy in comparison to separate analyses, a loss in prediction accuracy is expected relative to an analysis using all individual-level data due to approximations (Maier et al., 2018). Further, these methods are based on some assumptions that make them difficult to apply outside their context of development. For example, Maier et al. (2018) implicitly assumed that only a single phenotype record per trait was associated with an individual. While this is usually the case in human genetics, it is not in breeding populations where individuals may have repeated phenotype records for the same trait, e.g., repeated longitudinal production or reproduction records in livestock or replicated field trials in crops, or when phenotype records are measured on a group of individuals and linked to a genotyped relative, e.g., progeny tested bulls for dairy production.

The objective of this study was to develop a method that jointly analyses individual-level data and summary statistics from multiple populations with no or limited amount of approximation. The method assumes that individual-level data is composed of marker genotypes and phenotype records that potentially have a variable number of replicates per individual. Further, summary statistics are assumed to be composed of estimated allele substitution effects with an associated measure of accuracy. Different measures of accuracy can be used, which controls the amount of approximation. The developed method is validated with simulated data. The results show that the method enables accurate integration of different sources of information across a wide range of settings.

## MATERIAL AND METHODS

The first part of this section describes the theory of (1) separate and joint analyses of two individual-level datasets, (2) an exact integration of estimated allele substitution effects from one population into the analysis of another, (3) approximate integrations, and (4) generalization for multiple populations. The second part describes simulations used for validation of the developed method.

### Theory

Assume we have two populations with individual-level datasets of phenotyped and genotyped individuals. The two populations and their corresponding datasets are hereafter referred to as 1 and 2. Further assume that both datasets contain the same markers. From this data we want to obtain accurate estimates of allele substitution effects and genetic values for complex traits. We can achieve this by a joint analysis of the two datasets. When one of the datasets is not available, we can achieve this by integrating the results of a separate analysis of the unavailable data into the separate analysis of the available dataset. We show how to perform this integration exactly or approximately.

#### Separate and joint analyses

A standard marker model, using random regression on marker genotypes, for the separate analysis of dataset *i*(*i* = 1, 2) is:

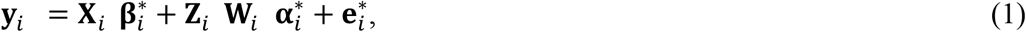

where **y**_*i*_ is a *n_obs,i_* × 1 vector of phenotypes, 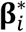 is a *n_f,i_* × 1 vector of fixed effects that are linked to **y**_*i*_ by a *n_obs,i_* × *n_f,i_* incidence matrix **X**_*i*_, 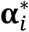 is a *n_mar_* × 1 vector of allele substitution effects that are linked to **y**_*i*_ by a *n_obs,i_* × *n_ind,i_* incidence matrix **Z**_*i*_ and a *n_ind,i_* × *n_mar_* matrix of genotypes **W**_*i*_, and 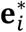 is the vector *n_obs,i_* × 1 of residuals. In this work we consider single-nucleotide polymorphism markers, which we code in **W**_*i*_ as 0 for homozygous aa, 1 for heterozygous aA or Aa, and 2 for homozygous AA. Other genotype coding and centering, that is of the form (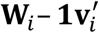) with **1** being a *n_ind,i_* × 1 vector of ones and **v**_*i*_ being a *n_mar_* × 1 vector, can be used with no difference in obtained estimates of allele substitution effects (Strandén and Christensen, 2011). We assume a prior multivariate normal (MVN) distribution for allele substitution effects for the separate analyis of the dataset *i*, 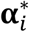 with mean zero and covariance 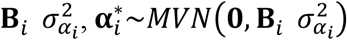 where **B**_*i*_ is a *n_mar_* × *n_mar_* diagonal matrix (e.g., an identity matrix **I**), and 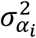 is the variance of allele substitution effects. We also assume that residuals are multivariate normally distributed with mean zero and covariance 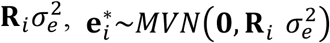, where **R**_*i*_ is a *n_obs,i_* × *n_obs,i_* diagonal matrix (e.g., an identity matrix **I**), and 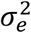 is the residual variance. For simplicity and without loss of generality, it is assumed in the following that residual variances are the same for all separate and joint analyses. Variance components 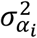 and 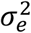 are assumed known, as they will have been estimated from the data previously. This marker model is the ridge regression model (Hoerl and Kennard, 1976; Whittaker et al., 2000; Meuwissen et al., 2001; de los Campos et al., 2012) with optional different weights in **B**_*i*_ (to differentially shrink different loci) and **R**_*i*_ (to account for heterogeneous residual variance due to variable number of repeated phenotype records per individual).

Separate estimates of allele substitution effects 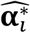 are obtained by solving the following system of equations:

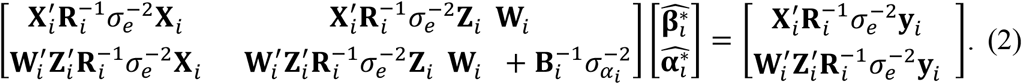

Separate estimates of genetic values for individuals in a dataset *i*(*i* = 1, 2) are obtained by 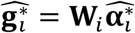.

A marker model for the joint analysis of two datasets 1 and 2 is:

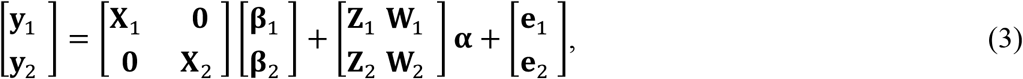

where phenotypes from the two populations are modelled with populations specific fixed effects (**β**_1_,**β**_2_) but a joint set of allele substitution effects (**α**) We assume a multivariate normal prior distribution for allele substitution effects with mean zero and covariance 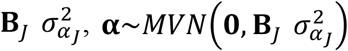, where **B**_*J*_ is a *n_mar_* × *n_mar_* diagonal matrix, and *σ_α_J__*^2^ is the variance of allele substitution effects in the joint analysis. We also assume that residuals are multivariate normally distributed, specifically 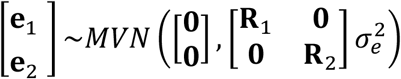 where **R**_*i*_ is a *n_obs,i_* × *n_obs,i_* diagonal matrix.

Joint estimates of allele substitution effects **α**̂ are obtained by solving the following system of equations:

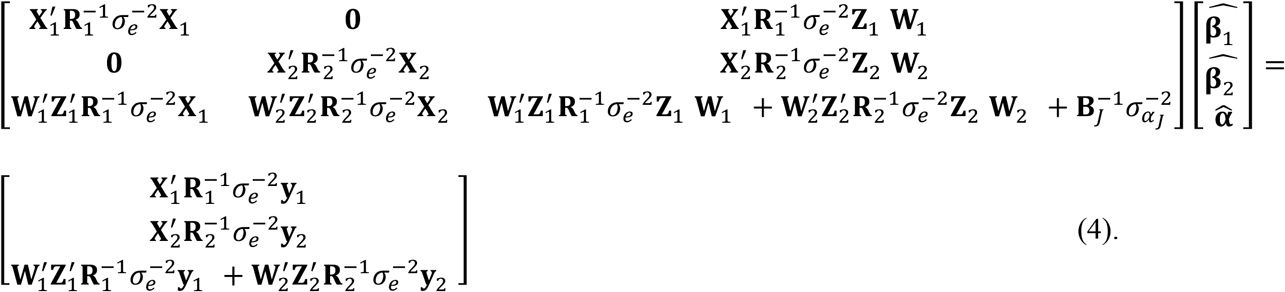

Joint estimates of genetic values for individuals in a dataset *i*(*i* = 1, 2) are obtained by 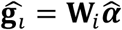.

#### Exact integration

The integration of estimates of allele substitution effects from one dataset into the analysis of another can be performed by means of absorbing corresponding equations in the joint system of equations. We choose to integrate estimates from the dataset 1 into the analysis of dataset 2. Derivations in Appendix A1 lead to the following system of equations that performs such integration and gives equivalent estimates of allele substitution effects to the joint analysis (4):

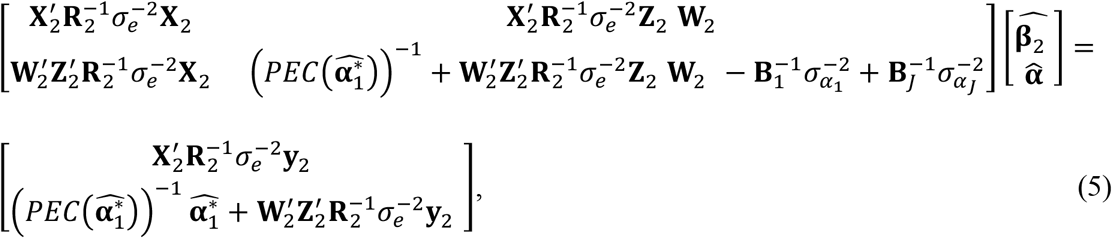

where 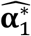 are estimates of allele substitution effects from the separate analysis of dataset 1 using (2), and 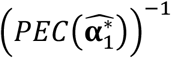 is the inverse of the corresponding prediction error covariance (PEC) matrix. The latter can be obtained as 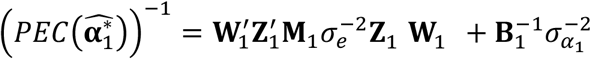 with 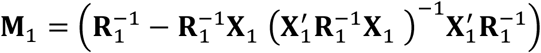. Note that only the individual-level dataset 2 and summary statistics from the dataset 1 (i.e., the estimated allele substitution effects and their PEC) are required. Individual-level dataset 1 is therefore not required.

It is worth noting that the integration of estimates of allele substitution effects from the dataset 1 into the analysis of dataset 2 can also be obtained from a Bayesian context. Bayes estimators for linear mixed models were discussed by several authors (Lindley and Smith, 1972; Dempfle, 1977; Gianola and Fernando, 1986). In a Bayesian context, we can assume the following prior multivariate normal distributions for the marker model (1) applied to dataset 2:

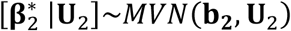, where **b**_2_ is a mean vector and **U**_2_ is a (co)variance matrix,
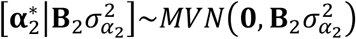, and
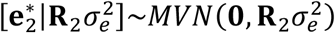

Assuming a noninformative prior for 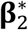 the system of equations (2) for dataset 2 can be obtained by differentiating the joint posterior distribution of 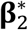 and 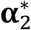 with respect to 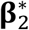 and 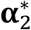, and setting the derivatives equal to 0 (Gianola and Fernando, 1986). Integration of estimates of allele substitution effects from dataset 1 into the analysis of dataset 2 can be therefore obained by defining a multivariate normal prior distribution for allele substitution effects in the analysis of dataset 2 using the posterior distribution for allele substitution effects from a separate analysis of dataset 1:

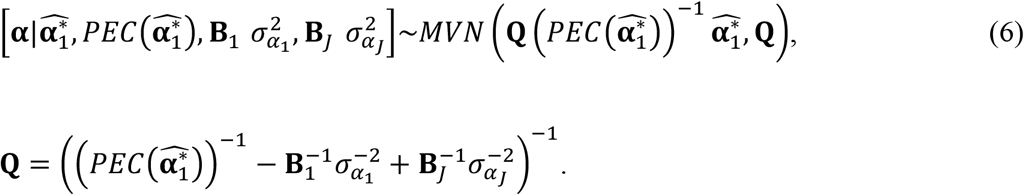

The matrix **Q** can be considered as the PEC matrix of a hypothetical separate analysis of dataset 1 using the multivariate normal prior distribution for allele substitution effects of the joint analysis, that is 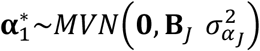 and 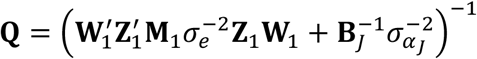, and the vector 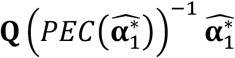 can be considered as the estimated allele subsitution effects of this hypothectical separate analysis. In animal breeding, a similar approach was used to integrate estimated genetic values and associated accuracies from one genetic evaluation into another genetic evaluation (Quaas and Zhang, 2006; Legarra et al., 2007; Vandenplas and Gengler, 2012).

Finally, it is worth noting that the term 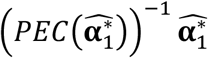 can be interpreted as pseudo-phenotypes associated with allele substitution effects of dataset 2, derived from information in dataset 1. In this sense, the system (5) is similar to approaches that compute pseudo-phenotypes from available estimated genetic values where individual-level phenotypic information is not readily available, or is not measured on the individuals themselves but on close relatives. In animal breeding, these approaches are commonly known as deregression of estimated genetic values (Jairath et al., 1998).

#### Approximate integration

Exact integration requires the inverse of prediction error covariance matrix from the separate analysis, which could be approximated when unavailable. Genomic analyses of complex traits that combine different datasets commonly have access to estimated allele substitution effects and associated prediction error variances (in different forms), but not the whole prediction error covariance matrix 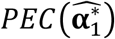 required in (5). We propose several ways to accommodate this situation. We assume that we know, at least, the prediction error variances (PEV) of estimated allele substitution effects (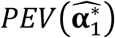), the number of individuals (*n*_*ind*,1_) and variance components used in the separate analysis of dataset 1 (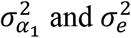).

When only the prediction error variances of the estimated allele substitution effects (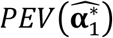) are known, while PEC are not, then we can approximate 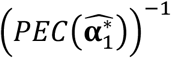 with 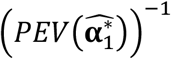. This approximation would be accurate if the matrix product 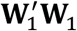 has (close to) zero off-diagonal elements, which is dependent on the characteristics of genotypes in dataset 1 (e.g., allele frequencies, linkage disequilibrium (LD), and population/family structure). If this is not the case, the approximation will bias the analysis by ignoring off-diagonal elements.

When allele frequencies and LD correlations in dataset 1 are known, we can obtain a good approximation of 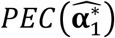 under some conditions (one phenotype record per individual, homogenous residual variance, overall mean is the only fixed effect, and Hardy-Weinberg equilibrium). Derivations in Appendix A2 show that under these conditions we can approximate 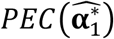 with 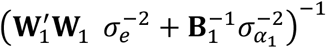 with the unknown matrix 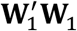 approximated from commonly available population parameters (i.e., allele frequencies and LD correlation) as 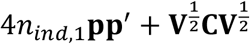, where **p** is a *n_mar_* × 1 vector of allele frequencies, **V** is a *n_mar_* × *n_mar_* diagonal matrix of expected genotype sum of squares with the *i*-th diagonal element equal to *n_ind_*_,1_ 2*p_i_*_,1_ (1–*p_i_*_,1_) and **C** is a *n_mar_* × *n_mar_* matrix of pairwise genotype correlations between markers. In practice, the matrix **C** for dataset 1 could be unknown, but we can approximate it by using a reference panel that includes, for example, available genotypes of non-phenotyped individuals originating from this population (Yang et al., 2012; Vilhjálmsson et al., 2015; Maier et al., 2018).

Finally, we relax the assumption of having a single phenotype record per individual in the preceding approximations. This is relevant when individuals have repeated phenotype records, e.g., repeated longitudinal production or reproduction records in livestock or replicated field trials in crops. A related issue is the violation of assumption of homogenous residual variance when phenotype records are first pre-processed and then used in genomic analyses, e.g., deregressed progeny proofs in livestock (e.g.,Garrick et al., 2009) or adjusted field trial means in crops (e.g.,Schulz-Streeck et al., 2013; Oakey et al., 2016; Damesa et al., 2017). For these situations, we show in Appendix A3 that we can approximate 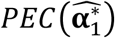 with 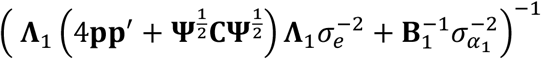 where **Ψ** is a *n_mar_* × *n_mar_* diagonal matrix with the *j*-th diagonal element equal to 2*p_j_*_,1_(1–*p_j_*_,1_) and **Λ**_1_ is a *n_mar_* × *n_mar_* diagonal matrix with the *j*-th diagonal element representing the square root of effective number of records for the *j*-th marker. The matrix **Λ**_1_ can be obtained by solving the nonlinear system of equations 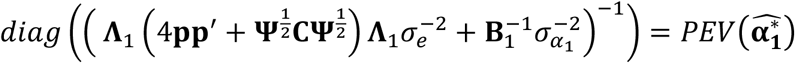 through a fixed-point iteration algorithm (Burden and Faires, 2010) detailed in Appendix A3. It is worth noting that the proposed algorithm requires the inversion of a *n_mar_* × *n_mar_* dense matrix at each iteration. This computational cost can be reduced by performing the algorithm for each chromosome separately.

#### Integration with multiple populations

When more than two populations or datasets are available the developed methods can be easily extended. With *n* datasets, the prior distribution for allele substitution effects in the separate analysis of the *n*-th dataset is defined using the posterior distributions for allele substitution effects from the separate analyses of *n*−1 datasets:

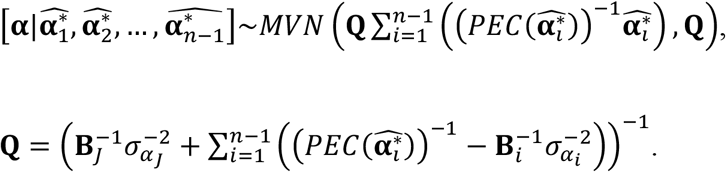

### Simulations

We tested developed methods with simulated data that either had low or high genetic diversity. The data was simulated in 5 replicates with the AlphaSim program, which uses the coalescent method for simulation of base population chromosomes and the gene drop method for simulation of chromosome inheritance within a pedigree (Hickey and Gorjanc, 2012; Faux et al., 2016).

A diploid genome was simulated with 30 chromosomes, each 108 base pairs long. Coalescent mutation and recombination rate per base pair were set to 10-8, while effective population size was modelled over time to mimic population history of a livestock population in line with the values reported by MacLeod et al. (2013). Specifically, for the low diversity scenario effective population size of the base population was set to 100 and increased to 120, 250, 350, 1,000, 1,500, 2,000, 2,500, 3,500, 7,000, 10,000, 17,000, and 62,000 at respectively 6, 12, 18, 24, 154, 454, 654, 1,754, 2,354, 3,354, 33,154, and 933,154 generations ago. For the high diversity scenario, effective population size of the base population was set to 10,000 and increased above this value in the same way as in the low diversity scenario; to 17,000 and 62,000 at 33,154, and 933,154 generations ago. For each chromosome 10,000 whole chromosome haplotypes were sampled, which on average hosted about 700,000 markers (21 million per genome) for the low diversity scenario and 1,400,000 markers (42 million per genome) for the high diversity scenario. Out of these loci 100 per chromosome (3,000 per genome) were sampled as causal loci affecting a complex trait. The allele substitution effect of causal loci was sampled from a normal distribution with mean zero and variance 1/3,000. The effects were used to simulate a complex trait with additive genetic architecture. In addition, 2,000 loci per chromosome (60,000 per genome) were selected as markers with the restriction of having minor allele frequency above 0.05.

From the base population, founder genomes for four populations (A, B, C, and D) were obtained by random sampling of chromosomes with recombination. The populations were ancestrally related through the common base population, but otherwise maintained independently, i.e., there was no migration between the four populations. Each population was initiated with 10,000 founders (half males and half females) and maintained for 7 generations with constant size. In the low diversity scenario, with the effective population size of 100, 25 males and 5,000 females were selected as parents of each generation, while in the high diversity scenario, with the effective population size of 10,000, all 5,000 males and 5,000 females were used. The 25 males were selected on true genetic value, assuming accurate progeny test was available.

For every individual in the population we simulated two types of phenotypes. First, an own single phenotype was simulated as the sum of the true genetic value and a residual sampled from a normal distribution with mean zero and residual variance scaled relative to the variance of true genetic value in the base population such that heritability was 0.3. These simulated single phenotype records mimic records measured on the individual. Second, a weighted phenotype was simulated as the sum of the true genetic value and the mean of *n_weight_* residuals. Each residual was sampled from a normal distribution with mean zero and residual variance scaled relative to the variance of true genetic value in the base population such that heritability was 0.3. The weight *n_weight_* was equal to *n_weight_* = 1+ *val* where the real value *val* was sampled from a geometric distribution with a probability of 0.15. The average *n_weight_* was 6.6. These weighted phenotypes mimic either repeated records of an individual or records on multiple progeny of an individual. To satisfy the assumption of identical residual variance across all analyses, phenotype records were divided by the residual standard deviation specific for each population, such that 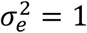. For every individual in each population we stored the true genetic value, own single and weighted phenotype records, associated weight, and 60,000 marker genotypes.

### Analysis

The data was analysed in several ways to evaluate the developed methods. In each case the aim was to obtain accurate genetic values utilizing all the available information. Specifically, we integrated results from separate analysis of populations B, C, and D, into the analysis of population A. We assumed throughout that variance components were known and equal to the rescaled variances. We analysed three scenarios in total. The first and second scenario used population specific training data of randomly sampled 30,000 individuals with single phenotype record from generations 1 to 6 under low and high diversity settings. The third scenario used population specific training data of randomly sampled 10,000 individuals with weighted phenotype record from generations 1 to 6 under low diversity setting. In all scenarios all of the 10,000 individuals from generation 7 of each population were considered as validation individuals. The following analyses were performed:

1) A joint analysis of four populations. This was the reference that the other analyses were compared against;
2) A separate analysis for each of the four populations;
3) An exact integration of separate analyses of populations B, C, and D, into the analysis of population A;
4) The same as 3), but approximating the PEC matrix with a partial PEC matrix for each chromosome, i.e., PEC between markers on different chromosomes were set to zero;
5) The same as 3), but approximating the PEC matrix with a diagonal PEV matrix, i.e., PEC between all markers were set to zero;
6) The same as 3), but approximating the PEC matrix with PEV, allele frequencies, and LD correlations between markers obtained from the training sets. For the scenario with weighted phenotype records, the algorithm for estimating the effective number of records per marker was performed for each marker separately and for each chromsome separately.
7) The same as 6), but with LD correlations between markers computed from validation individuals instead of the training data.

For each analysis we calculated genomic prediction accuracy as the Pearson correlation between the true and estimated genetic value in validation individuals. Further, we evaluated the different integrations by comparing estimated genetic values of validation individuals against the estimated genetic values obtained from the joint analysis, which was considered as the reference because it used information from all populations. If integration was fully accurate, there should be no difference between the joint analysis and the analysis with integration. We assessed this by (a) accuracy of integration as a Pearson correlation between estimated genetic values from the joint analysis and the analysis with integration (desired value equals 1), (b) calibration of integration as a regression of estimated genetic values from the joint analysis on estimated genetic valuesfrom analysis with integration (desired value equals 1), and (c) magnitude of error in integration as a mean square error (MSE) between estimated genetic values from the joint analysis and from the analysis with integration (desired value equals 0).

### Data availability

Supplemental figures are available in File S1. A description of the simulated genotype and phenotype datasets for each scenario is provided in File S2. Simulated genotype and phenotype datasets for the 5 replicates of each scenario are provided in Files S3, S4, and S5. All files were uploaded to Figshare.

## RESULTS

### Genomic prediction accuracy of separate and joint analyses

Joint analysis increased genomic prediction accuracy in comparison to separate analyses. This is shown in Table 1. Analysing separately the four datasets gave accuracies of about 0.71 (low diversity) and 0.53 (high diversity) with single phenotype records, and of about 0.73 (low diversity) with weighted phenotype records. Analysing jointly the four datasets increased accuracy by 0.09 absolute points with single phenotype records and by 0.12 absolute points with weighted phenotype records.

**Table 1.** Genomic prediction accuracy for joint and separate analyses in scenarios with single or weighted phenotype records and low or high diversity (values are averages across the five replicates^1^)

| Phenotypes | Diversity | Analysis | Populations |  |  |  |
| --- | --- | --- | --- | --- | --- | --- |
|  |  |  | A | B | C | D |
| Single | Low | Joint | 0.811 | 0.811 | 0.823 | 0.815 |
|  |  | Separate | 0.705 | 0.708 | 0.718 | 0.718 |
|  | High | Joint | 0.687 | 0.686 | 0.687 | 0.684 |
|  |  | Separate | 0.536 | 0.537 | 0.528 | 0.528 |
| Weighted | Low | Joint | 0.860 | 0.865 | 0.865 | 0.862 |
|  |  | Separate | 0.720 | 0.739 | 0.724 | 0.727 |

<sup>1</sup> Standard errors are between 0.003 and 0.016.

### Integration based on PEC, partial PEC, or PEV matrices

For all scenarios the developed method enabled exact integration when complete PEC matrices were used. Integration of estimated allele substitution effects by means of the complete PEC matrix led to the same estimated genetic values as with the joint analysis, as shown by correlation and regression coefficients of 1, and MSE close to 0 (Figures 1-6; Figures S1-S6). For comparison, correlations between estimated genetic values from separate analyses and joint estimated genetic values were about 0.87 (low diversity) and 0.77 (high diversity) with single phenotype records, and 0.85 (low diversity) with weighted phenotype records.

**Figure 1.**
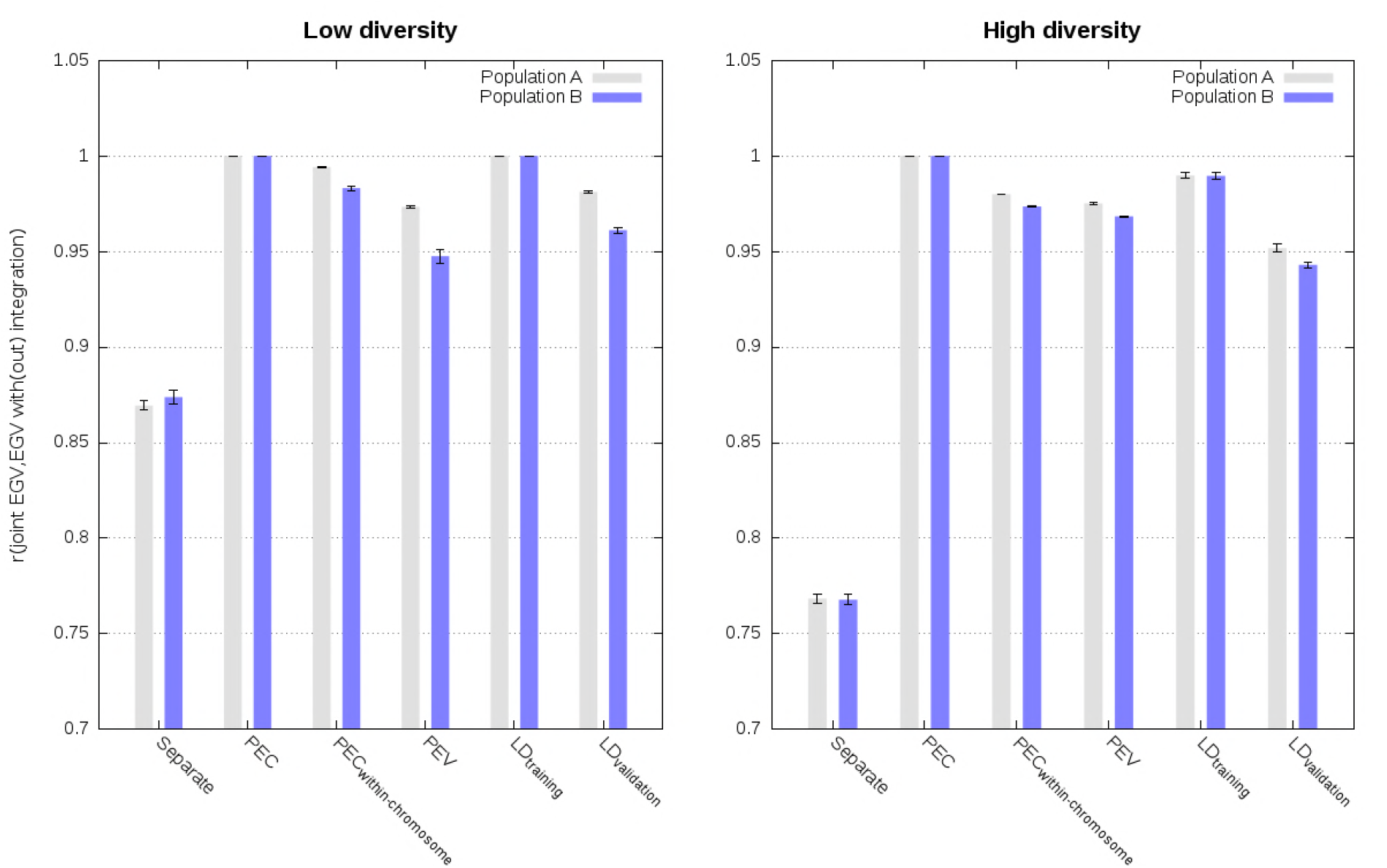
Correlation between estimated genetic values (EGV) from the joint analysis and from different analyses in populations A and B using a single phenotype record per individual in scenarios with low and high diversity (values are averages across the five replicates with standard errors).

Approximate integration by means of partial PEC matrices for each chromosome, that is ignoring PEC between markers on different chromosomes, gave almost as accurate and calibrated estimated genetic values as the exact integration. This is illustrated in Figures 1-6 with correlations higher than 0.96, regression coefficients close to 1, and MSE close to 0. Increasing the diversity slightly deteriorated accuracy and calibration of genomic predictions (Figures 1-3; Figures S1-S3).

Approximate integrations by means of PEV matrices, that is ignoring PEC between all markers, gave quite accurate, but uncalibrated estimated genetic values. This is shown in Figures 1-6 and in Figures S1-S6. Correlations between joint estimated genetic values and estimated genetic values with integration by means of PEV were between 0.95 and 0.98 with single phenotype records and between 0.93 and 0.95 with weighted phenotype records Despite these correlations close to 1, estimated genetic values were uncalibrated, as depicted by regression coefficients below 0.77 for the low diversity scenarios with single and weighted phenotype records, and below 0.86 for the high diversity scenario with single phenotype records (Figures 2, 5, S2, S5).

**Figure 2.**
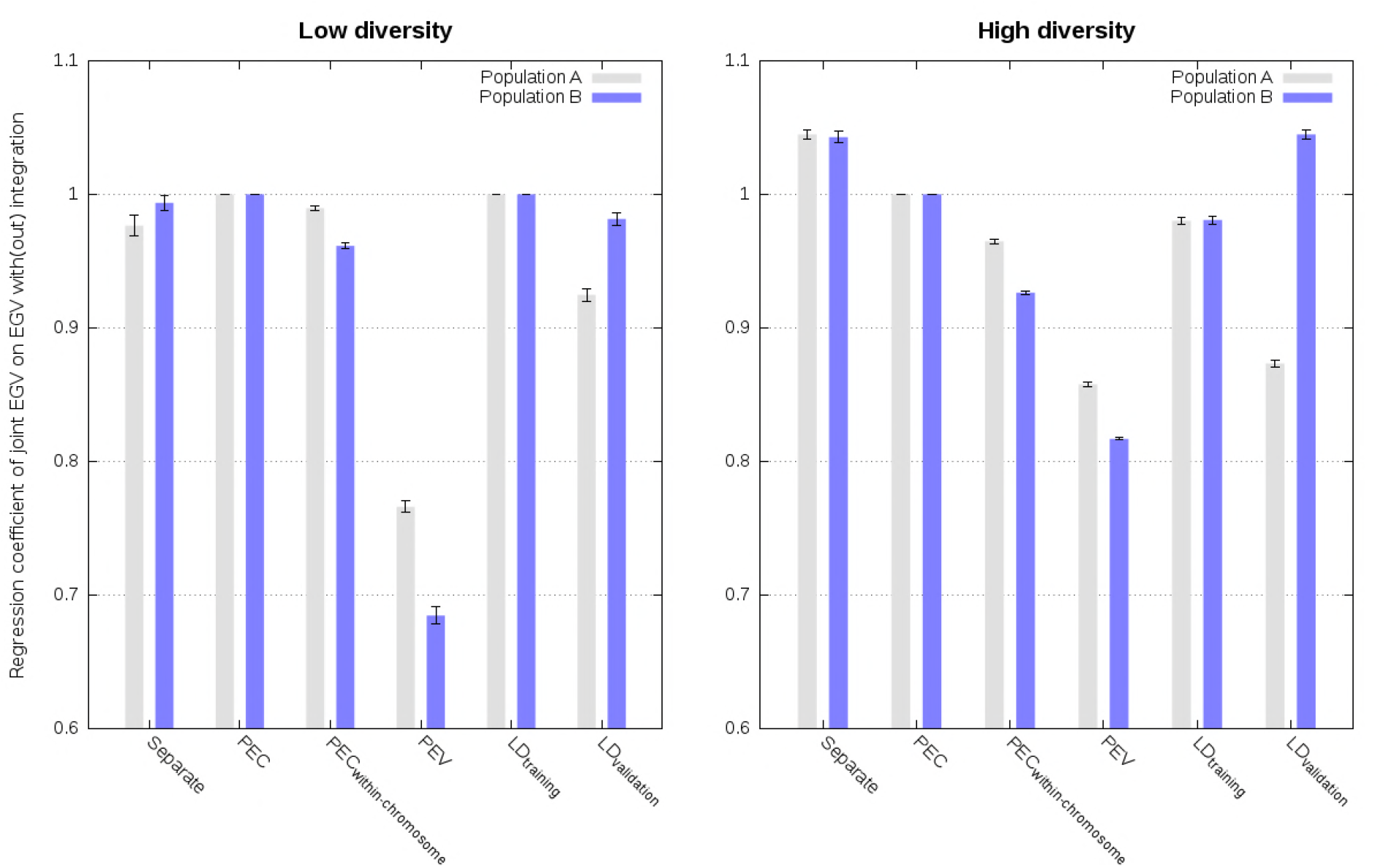
Regression of estimated genetic values (EGV) from the joint analysis on estimated genetic values from different analyses in populations A and B using a single phenotype record per individual in scenarios with low and high diversity (values are averages across the five replicates with standard errors).

### Integration based on PEV, allele frequencies, and LD information

When LD information was derived from training data of other populations, approximate integrations by means of PEV, allele frequencies, and LD information, resulted in highly accurate and well calibrated estimated genetic values with single phenotype records. This is shown in Figures 1-3 (Figures S1-S3). Correlation and regression coefficients were equal to 1 for the low diversity scenario. Slightly lower values, but still close to 1, were observed for the high diversity scenario. For both low and high diversity scenarios, MSE were close to 0. In contrast, when LD information was derived from validation data of other populations, approximate integrations gave less accurate and well calibrated estimated genetic values. This is shown in Figures 3-6 (Figures S3-S6). For these scenarios, correlations were equal to at least 0.94, and regression coefficients varied between 0.87 and 1.05.

**Figure 3.**
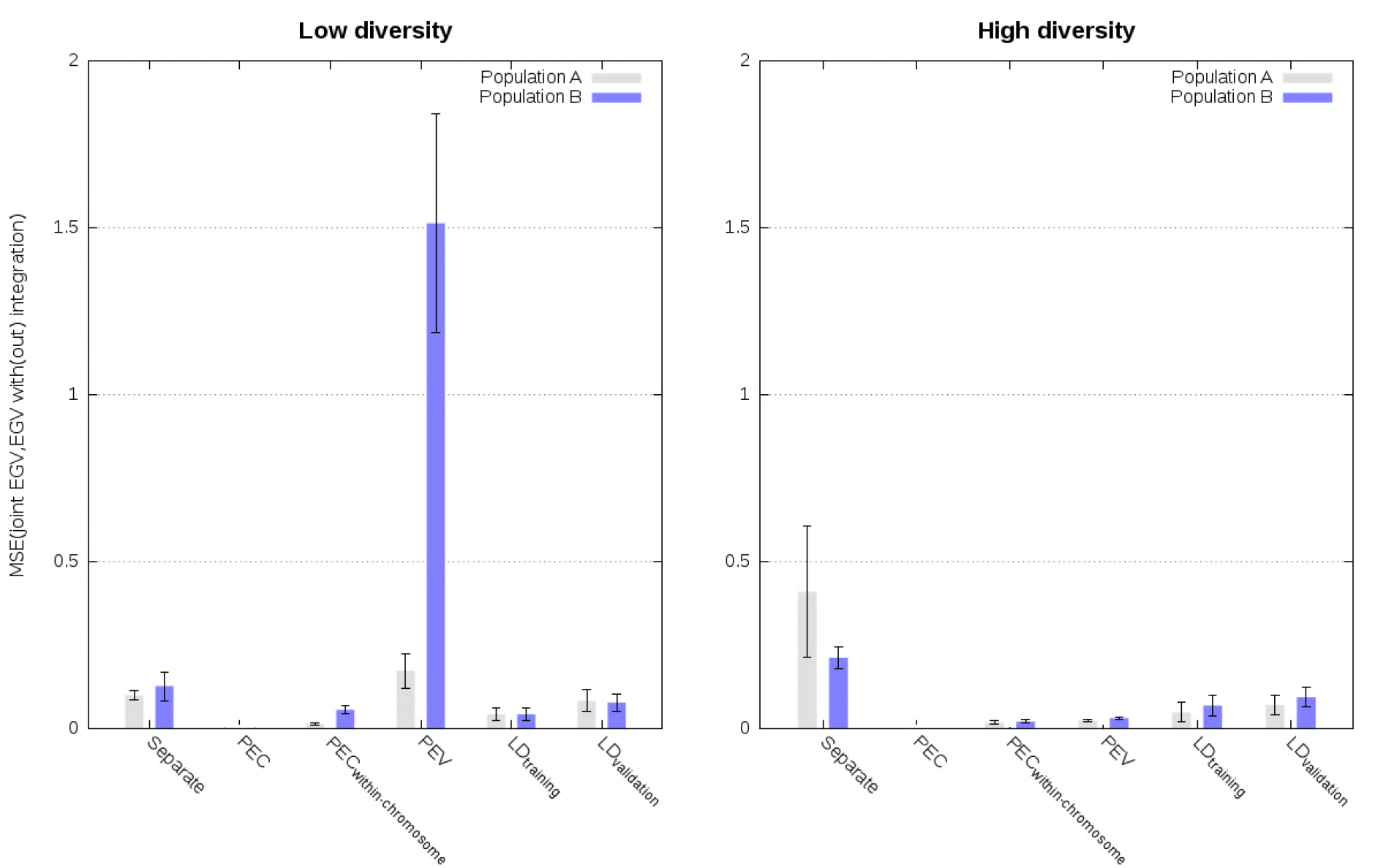
Mean square errors between joint estimated genetic values (EGV) from the joint analysis and from different analyses in populations A and B using a single phenotype record per individual in scenarios with low and high diversity (values are averages across the five replicates with standard errors).

For the scenario with weighted phenotype records, approximate integrations by means of LD information from training data of other populations resulted in highly accurate and well calibrated estimated genetic values when sets of markers per chromosome were used to estimate the effective number of records for each marker. Correlations between joint estimated genetic values and estimated genetic values with integration were about 0.99 (Figure 4, Figure S4), regression coefficients were about 0.95 (Figure 5, Figure S5), and MSE were close to 0 (Figure 6, Figure S6). Using LD information from the validation data of other populations, instead from the training data of other populations, gave slightly less accurate (correlations higher than 0.95), and moderately less calibrated estimated genetic values (regression coefficients between 0.87 and 1.04; Figure 4-6; Figures S4-S6). For both cases, estimating the effective numbers of records per marker, instead of for all markers per chromosome simultaneously, reduced accuracy and calibration of estimated genetic values (Figure 4-5; Figures S4-S5).

**Figure 4.**
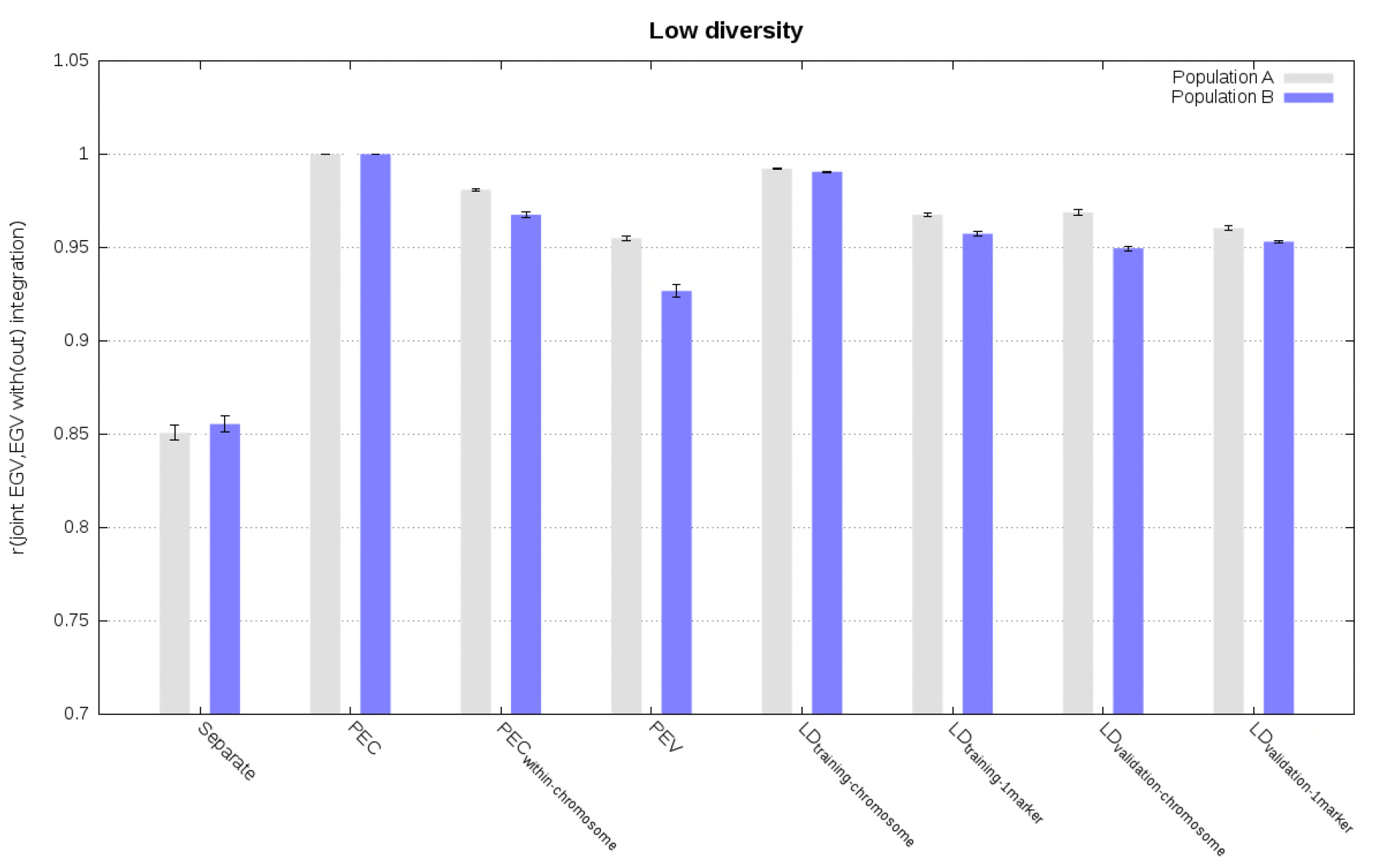
Correlation between estimated genetic values (EGV) from the joint analysis and from different analyses in populations A and B using weighted phenotype records in the scenario with low diversity (values are averages across the five replicates with standard errors).

**Figure 5.**
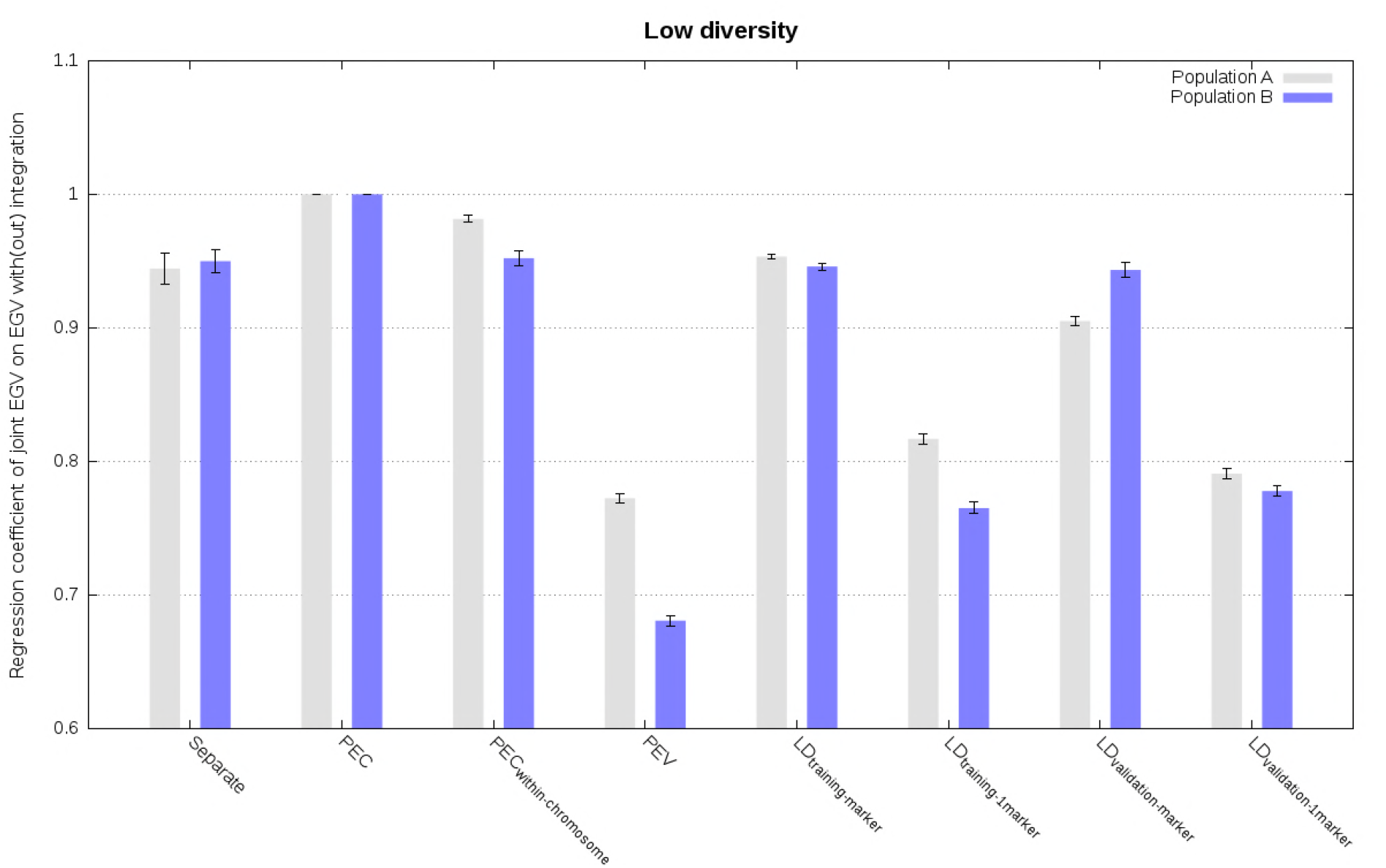
Regression of estimated genetic values (EGV) from the joint analysis on estimated genetic values from different analyses in populations A and B using weighted phenotype records in the scenario with low diversity (values are averages across the five replicates with standard errors).

**Figure 6.**
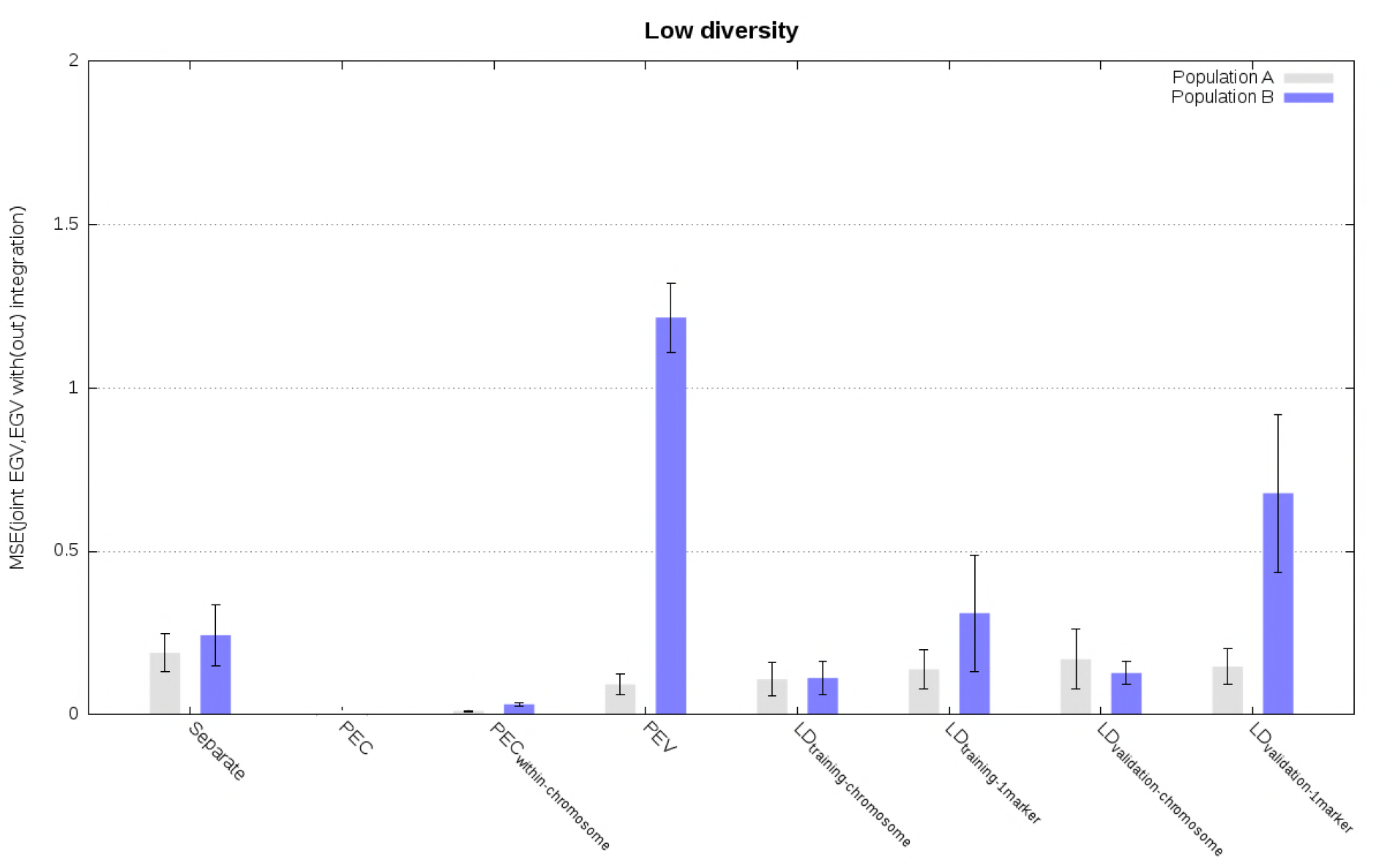
Mean square errors (SE) between estimated genetic values (EGV) from the joint analysis and from different analyses in populations A and B using weighted phenotype records in the scenario with low diversity (values are averages across the five replicates with standard errors).

### Comparison of estimated allele substitution effects

Correlation and regression coefficients between estimated allele substitution effects from the joint analysis and analysis with integration largely followed patterns of the corresponding values for estimated genetic values (Tables 2-3). Correlation and regression coefficients were close to 1 when the integration of estimated allele substitution effects was by means of the complete PEC matrices. Ignoring PEC between markers on different chromosomes, or ignoring PEC between all markers, reduced correlations to between 0.92 and 0.99 (Tables 2-3). Using LD information with PEV led to correlations between joint estimates of allele substitution effects and estimates with integration ranging from 0.71 to 0.83 for the scenario with weighted phenotype records (Tables 2-3).

**Table 2.** Comparison of estimated allele substitution effects from different analyses with estimates from the joint statistical analysis using single phenotype records in scenarios with low and high diversity (values are averages across the five replicates^1^)

| Analysis | Low diversity |  | High diversity |  |
| --- | --- | --- | --- | --- |
|  | Correlation | Regression | Correlation | Regression |
| Separate A | 0.71 | 1.09 | 0.65 | 1.10 |
| Separate B | 0.71 | 1.09 | 0.65 | 1.10 |
| Separate C | 0.71 | 1.09 | 0.65 | 1.11 |
| Separate D | 0.71 | 1.09 | 0.64 | 1.10 |
| PEC | 1.00 | 1.00 | 1.00 | 1.00 |
| PEC<sub>within chromosome</sub> | 0.99 | 0.98 | 0.97 | 0.95 |
| PEV | 0.96 | 0.80 | 0.96 | 0.89 |
| LD<sub>training</sub> | 1.00 | 1.00 | 0.98 | 0.97 |
| LD<sub>validation</sub> | 0.96 | 0.88 | 0.93 | 0.84 |

<sup>1</sup> Standard errors are between 0.00 and 0.01.

**Table 3.**
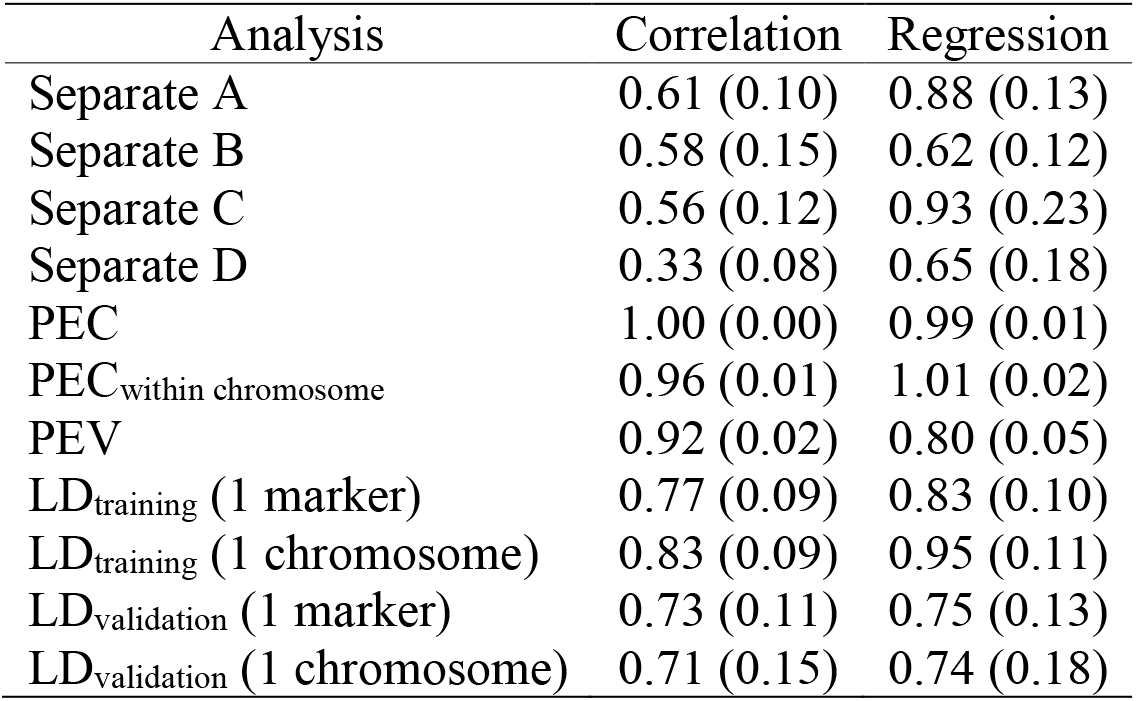
Comparison of estimated allele substitution effects from different analyses with estimates from the joint statistical analysis using weighted phenotype records in the scenario with low diversity (values are averages across the five replicates with standard errors between brackets)

## DISCUSSION

The results show that the developed method enables accurate and well calibrated estimated genetic values for complex traits using both individual-level data and summary statistics. As expected from theory, the analysis of individual-level data and estimated allele substitution effects from other analyses by means of PEC matrices, yielded the same estimates as the joint analysis of all individual-level data. To our knowledge, this is the first time that individual-level data and summary statistics were analysed simultaneously for genomic predictions. As illustrated by simulations, the combined analysis of multiple datasets may increase genomic prediction accuracy over separate analyses of a single dataset. Unfortunately, combining individual-level data from several sources is generally not feasible for several reasons, e.g., political roadblocks, data protections concerns, or data inconsistencies (Powell and Sieber, 1992; Vilhjálmsson et al., 2015; Maier et al., 2018). However, summary statistics, such as estimates of allele substitution effects and associated measures of accuracy (e.g., PEV), are usually available for exchange. The developed method enables increase in genomic prediction accuracy of complex traits by means of jointly analysing the available individual-level data and summary statistics.

Accurate integration of estimated allele substitution effects is possible also when the complete PEC matrix is not available. This is important because computing the exact PEC matrix and exchanging it between analyses might be challenging in some cases. For the vast majority of used marker arrays in animal and plant breeding the calculations and data transfers should be doable. For example, most arrays have between 10,000 and 100,000 markers, for which we need between ~1 and ~80 GB of memory to store the PEC matrix and between a minute and a day to invert it on current computers. For a larger number of markers, commonly used in human genetics, the memory requirements and computing time become prohibitive. The results show that in such cases we can still obtain accurate genomic predictions when the integration is done by means of partial PEC matrices for each chromosome. This is expected since high LD between markers mostly occurs within chromosomes. High LD between markers on different chromosomes may especially occur in structured populations and populations under selection (Farnir et al., 2000; Flint-Garcia et al., 2003; Rostoks et al., 2006). Both of these conditions are present in breeding populations. However, the results suggest that LD between chromosomes can be ignored for the purpose of integration for populations with both low and high diversity. The results also show that we can succesfully integrate estimated allele substitution effects when only PEV and allele frequencies from each population are available together with LD information of a reference genotype panel representative of each population. Assuming that such reference genotype panels are available, only estimated allele substitution effects, associated PEV, and allele frequencies need to be exchanged between populations for such analyses. Similar conclusions were drawn from studies combining only summary statistics obtained from genome-wide association studies to perform multi-trait genomic predictions (Maier et al., 2018).

Accurate integration of estimated allele substitution effects is possible irrespective of the diversity of the populations and characteristics of genotypes (e.g., allele frequencies, LD). This is obvious, and confirmed by our results, when integration is perfomed by means of complete PEC matrices. When complete PEC matrices are unavailable, accurate integration is possible if the inverses of the PEC matrices can be approximated accurately from available population parameters (i.e. LD and allele frequency information), whatever the level of diversity and characteristics of the populations, as shown by our results or a study combining summary statistics in human genetics (Maier et al., 2018). In our study, the population parameters obtained from the reference panels adequately reflected the characteristics of the training sets. Future studies should be conducted to assess the impact of suboptimal reference panels. Therefore, the developed method is expected to perform well on any type of data, from animal and plant breeding to human genetics, provided accurate information is available.

The developed method has some simplifying assumptions that can be readily relaxed. For example, we assumed that the same genotype coding was used in all populations. This assumption can be relaxed when centered genotype coding (i.e., of the form of (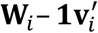)) is used because variance component estimates, estimates of allele substitution effects and PEC are the same irrespective of the centering of the genotype coding, provided that the model has a fixed general mean, which is considered in the integration (Strandén and Christensen, 2011). Also, centered and scaled (standardised) genotype coding is often used in human genetics, instead of only centered genotype coding (Yang et al., 2010; Speed et al., 2012; Maier et al., 2018). In practice, estimated genetic values are not influenced by scaling of centered genotype coding (Strandén and Christensen, 2011; Bouwman et al., 2017). Therefore, allele substitution effects estimated using one type of genotype scaling could be obtained from a post-analysis by converting estimated genetic values computed for a reference genotype panel into allele substitution effects for another genotype scaling. Converting estimated genetic values into allele substitution effects is often referred to as back-solving of allele substitution effects (Strandén and Garrick, 2009; Strandén and Christensen, 2011; Wang et al., 2012; Bouwman et al., 2017). Prediction error covariances associated with the converted estimated allele subsitution effects could be derived from the (prediction error) covariances of the estimated genetic values (see derivations in Appendix A4).

Allele substitution effects estimated from analyses using different different sets of markers or different residual variances, can be used in the integration as well. The assumption that all individuals were genotyped at the same loci could be considered as fullfilled if small differences in the sets of markers are corrected by assuming zero allele substitution effect and zero accuracy for markers not used in an analysis. When large differences between sets of markers are observed, this assumption can be accomodated following two approaches. A first, post-analysis, approach consists of assuming that estimated genetic values are the same for two different sets of markers, allowing the conversion of estimated allele substitution effects from one set of markers to another set of markers (Liu and Goddard, 2018). The conversion can be performed by back-solving estimated allele substitution effects from estimated genetic values, as proposed previously for different genotype codings, or by applying a marker model to the estimated genetic values with the reference set of markers (Liu and Goddard, 2018). A second approach consists of harmonizing genotype data across populations. This approach must be performed before the analyses, and requires therefore coordination between populations. Harmonization of genotype data could be performed by identifying a subset of markers for which all populations are genotyped, or by genotype imputation (e.g.,Marchini and Howie, 2010). Finally, the assumption that residual variances were the same in all populations, can be relaxed by noting that separate estimates of allele substitution effects 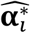, obtained by the system of equations (2), can be also obtained by the following different formulations:

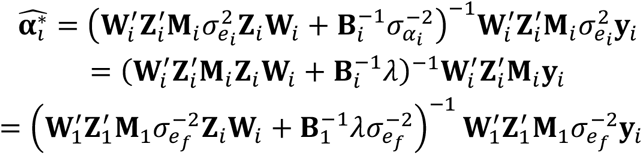

where 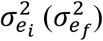 is the residual variance used for the *i*-th (focal) analysis, and 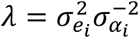.

For integration of 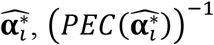 must be approximated using the residual variance of the focal population (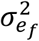) and the effective numbers of records per marker estimated using variance components of the *i*-th analysis. Another way to relax this assumption is to extend our univariate model to a bivariate model, similarly to methods developed to combine different genetic evaluations in animal breeding (Schaeffer, 1994; Vandenplas et al., 2015). In a bivariate model, one trait would represent individual-level data, while the other trait would represent summary statistics. The genetic correlation between the two traits could be estimated based on a subset of individual-level data available for both datasets or based on summary statistics (Bulik-Sullivan et al., 2015). Such an approach would also allow the integegration of summary statistics expressed on a different scale (e.g., different measure units, trait definitions) than the scale of the focal population (Vandenplas et al., 2015).

The developed method can be readily generalized to multi-trait models and is therefore a generalization of previous works that were based on several (implicit) assumptions (Liu and Goddard, 2018; Maier et al., 2018). For example, previous works assumed that no individual-level data were available. It was also (implicitly) assumed that only single phenotype records with homogeneous residual variance (Maier et al., 2018), or that the least-squares part of the separate analyses (Liu and Goddard, 2018), were available for integrating estimated allele substitution effects. Both assumptions lead to simple and accurate approximations of PEC matrices as shown in our study. However, we relax all these assumptions, such that our method can jointly analyse individual-level data and summary statistics, with possibly multiple phenotype records per individual.

## CONCLUSIONS

We developed a method for genomic prediction that accurately integrates summary statistics obtained from analyses of separate populations into an analysis of individual-level data. The method accommodates use of multiple phenotype (pseudo-)records per individual, and further extensions have been presented to accommodate for differences in residual variances or genotype codings used in the populations. When complete summary statistics information is available the method gives identical genomic predictions as the joint analysis of individual-level data from all populations. When summary statistics information is not complete we can use a series of approximations that give very accurate and well calibrated genomic predictions.

## ACKNOWLEDGMENTS

This study was financially supported by the Dutch Ministry of Economic Affairs (TKI Agri & Food project 16022), the Breed4Food partners Cobb Europe, CRV, Hendrix Genetics and Topigs Norsvin, and UK Biotechnology and Biological Sciences Research Council (BBSRC) ISPG to The Roslin Institute BBS/E/D/30002275. The use of the HPC cluster has been made possible by CAT-AgroFood (Shared Research Facilities Wageningen UR).

## Appendix A1: Exact integration

Here we detail the derivation of exact integration by means of absorbing the set of equations that pertain to one dataset. We start with the system of equations for separate analysis of dataset 1:

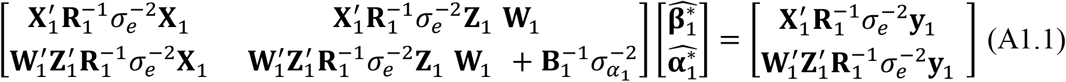

and the system of equations for the joint analysis of datasets 1 and 2:

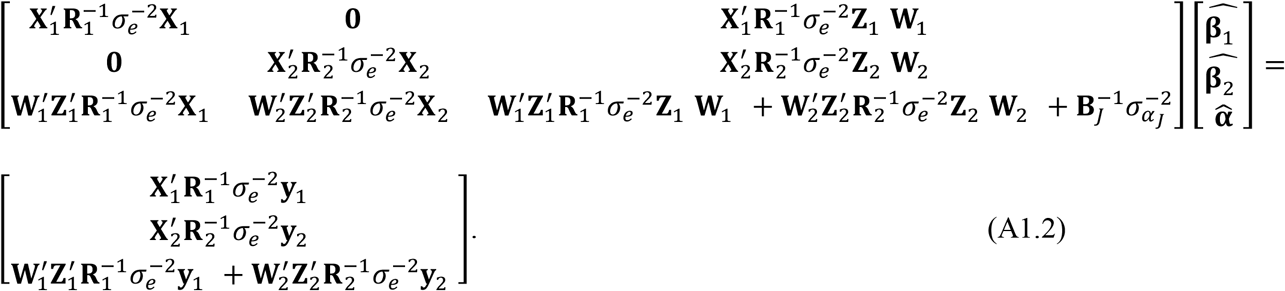

From the first set of equations 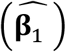 in (A1.2) it follows:

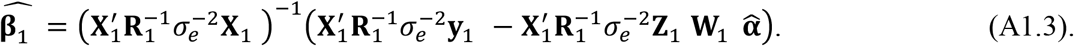

From the third set of equations 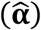 in (A1.2) it follows:

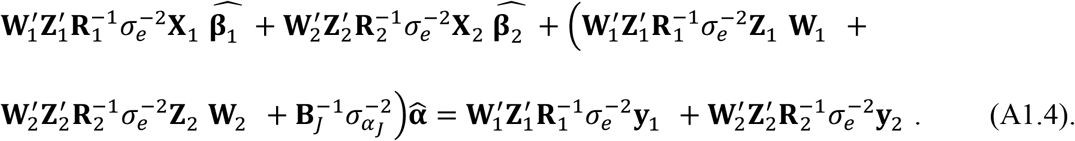

Inserting (A1.3) into (A1.4) gives, after some algebra:

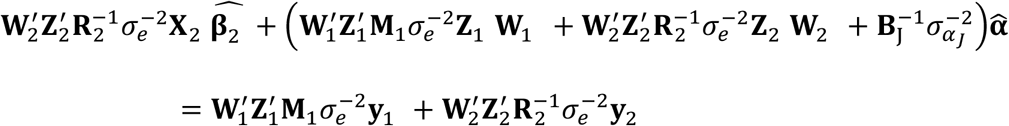

with 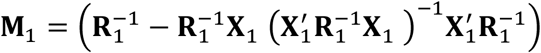

Now the system of equations (A1.2) can be re-written with the first set of equations (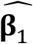) absorbed as:

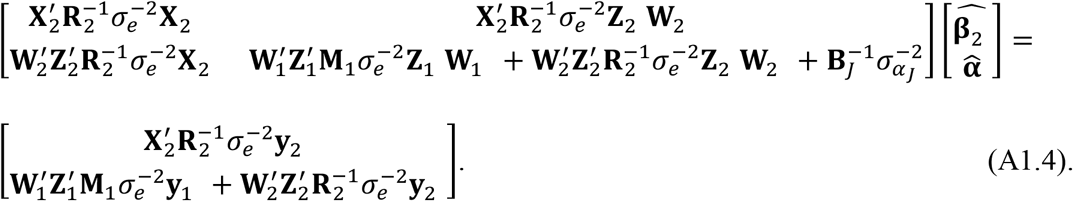

Similarly, the absorption of the first set of equations (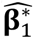) in separate analysis of dataset 1 (A1.1) leads to

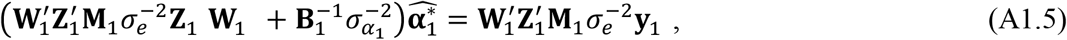

where

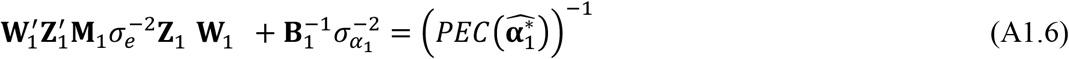

is the inverse matrix of prediction error covariances of 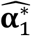.

Combining (A1.4) and (A1.5) with the use of (A1.6) enables the exact integration of estimates from the separate analysis of dataset 1 into the separate analysis of dataset 2 with the following system of equations:

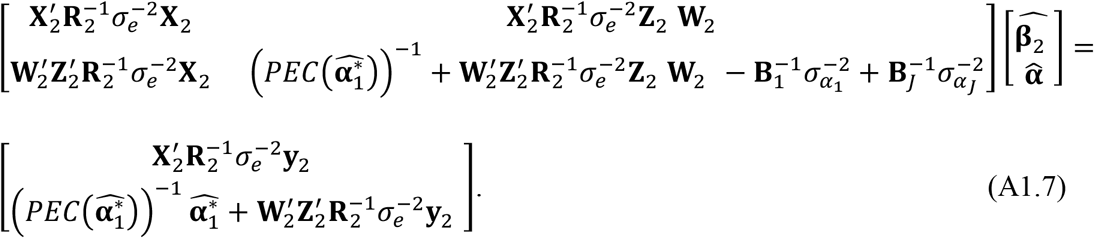

## Appendix A2: Approximate integration

Here we detail the derivation of different approximate integrations by means of simplified assumptions and use of summary statistics. We start with the expression for prediction error covariance matrix of allele substitution effects from dataset 1:

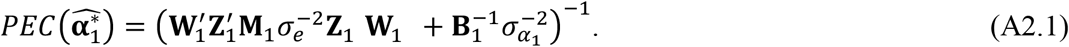

If we assume that: (1) every individual has a single phenotype record, i.e., **Z**_1_ = **I**,(2) residual variance is homogeneous, i.e. **R**_1_ = **I**, and (3) only overall mean is fitted as a fixed effect, i.e., **X**_1_ = **1**; then we can simplify (A2.1) as:

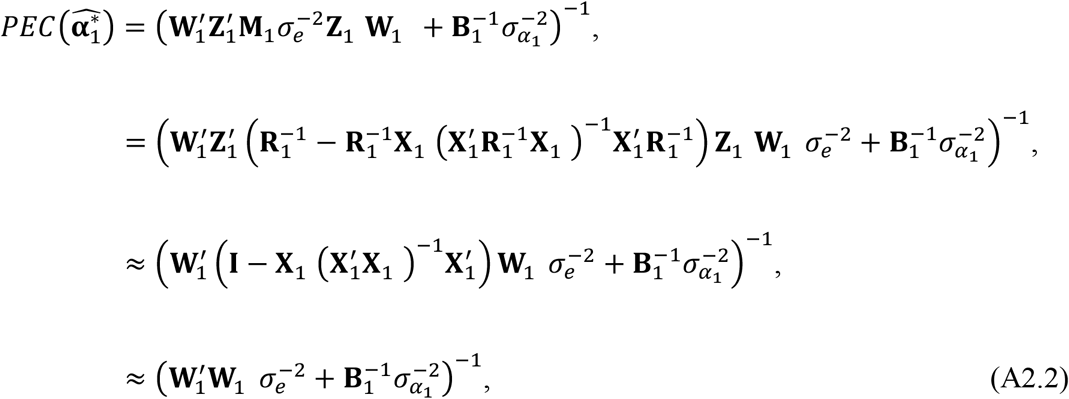

because 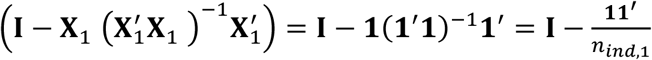 will tend to the identity matrix **I** with increasing *n_ind_*_,1_. The matrix (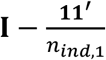), also known as the centering matrix, is a symmetric and idempotent matrix with off-diagonal elements equal to 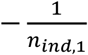 and with diagonal elements equal to 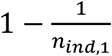.

When genotypes from the dataset 1 are not available, but variance components 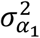 and 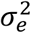 are, we “only” need to approximate the unknown matrix of genotype sum of squares 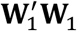 in (A2.2). This product can be approximated from linkage-disequilibrium and allele frequency information of the dataset 1, as shown in the following (similarly to Yang et al. (2012), Vilhjálmsson et al. (2015), and Maier et al. (2018)). Assume that linkage-disequilibrium between two markers is represented by the correlation of their unphased genotypes (Rogers and Huff, 2009). Then, a matrix of all pairwise correlations between markers is:

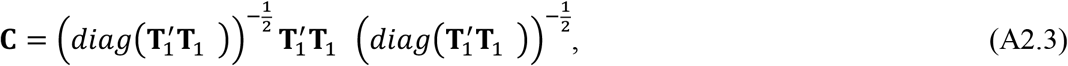

where the matrix **T**_1_ contains centered genotypes of dataset 1 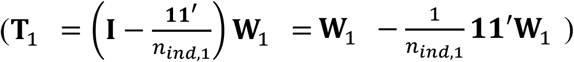. The matrix product 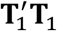 can be computed as:

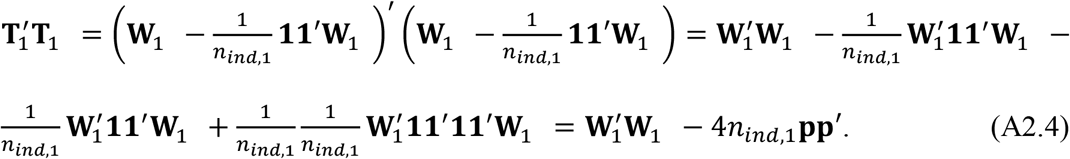

where 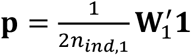 are allele frequencies in dataset 1 (Strandén and Christensen, 2011). Assuming Hardy-Weinberg equilibrium, the *i*-th diagonal element of the matrix product 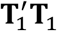, is equivalent to expected genotype sum of squares at the *i-*th marker, *n_ind_*,_1_ 2*p_i_*_,1_(1 − *p_i_*_,1_) with *p_i_*_,1_ being the allele frequency of the *i*- th marker in dataset 1.

Combining (A2.3) and (A2.4) we can approximate the unknown matrix of genotype sum of squares 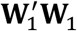 as:

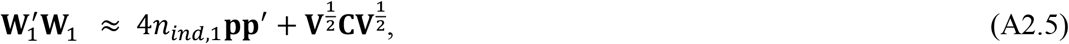

where **V** is diagonal matrix of expected genotype sum of squares with the *i*-th diagonal element equal to *n*_*ind*,1_2*p*_*i*,1_(1 − *p*_*i*,1_).

## Appendix A3: Estimation of the effective number of records per marker

Here we detail the algorithm for computing the effective number of records per marker by use of available population parameters (i.e. linkage-disequilibrium, and allele frequency information) and prediction error variances of 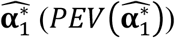 of the dataset 1. We start with the expression for the prediction error covariance matrix of allele substitution effects from dataset 1:

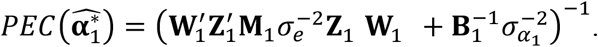

If the number of individuals and the number of records per individual are unknown, we can assume that a *n_mar_* × *n_mar_* diagonal matrix **Λ**_1_ exists such that:

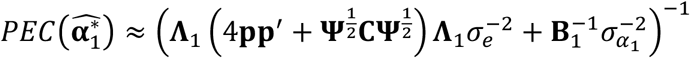

where **Ψ** is a *n_mar_* × *n_mar_* diagonal matrix with the *j*-th diagonal element equal to 2*p_j_*_,1_(1 − *p_j_*_,1_), and the squared *j-* th diagonal element of **Λ**_1_ represents the effective number of records for the *j*-th marker. The term (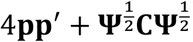) is similar to the approximation of the unknown matrix of genotype sum of squares 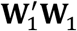 (i.e., 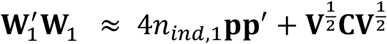) in the Appendix A.2. However, it does not involve the number of individuals *n_ind_*_,1_ because it is confounded with the effective number of records.

The diagonal matrix **Λ**_1_ can be estimated by solving the nonlinear system of equations 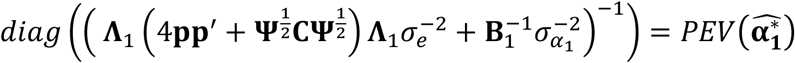 through a fixed-point iteration algorithm (Burden and Faires, 2010) as follows:

1. 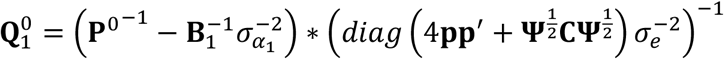 Where **p**^0^is a diagonal matrix with the *i*-th diagonal element equal to the PEV of the *i*-th marker and 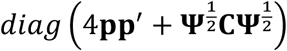 contains the diagonal elements of (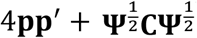);
2. 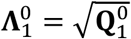
3. *k* = 1
4. 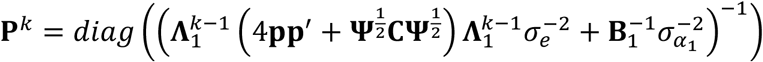
5. 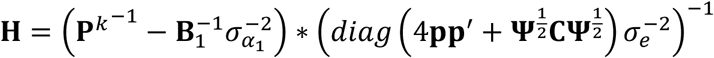
6. 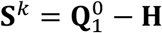
7. If trace of **S**^*k*^ is not sufficiently small:

a. 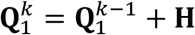
b. If any diagonal element in **Q**_1_^*k*^ is negative, set it to 0
c. 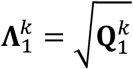
d. *k* = *k* + 1
e. Repeat from 4
8. 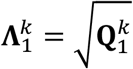

It is worth noting that the proposed algorithm is similar to algorithms to estimate effective number of records per individual, where “effective” means that they are free of contributions from relatives (Misztal and Wiggans, 1988; Vandenplas and Gengler, 2012). The *j*-th diagonal element of 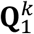 can therefore equivalently be considered as the effective number of records for the *j*-th marker.

## Appendix A4: Conversion of allele substitution effects

Here we detail a post-analysis to obtain allele substitution effects estimated using one type of genotype coding (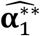) by converting estimated genetic values computed for a reference genotype panel with allele substitution effects for another genotype coding (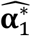) We assume that allele substitution effects (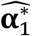) are available with the associated prediction error (co)variance matrix (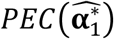), as well as the (co)variance matrix of 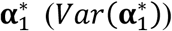, and genotypes of a reference panel using a particular type of genotype coding (**Γ**^*^)Estimates of genetic values for the reference individuals are obtained as 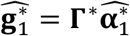.

Assuming that estimated genetic values are not influenced by scaling of centered genotype coding (Strandén and Christensen, 2011; Bouwman et al., 2017), and that the (co)variances of genetic values are the same irrespective of the genotype coding, we can write that 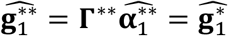 being a matrix with reference genotypes using another type of genotype coding than **Γ**^*^ and 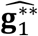 being a vector of estimated genetic values using this type of genotype coding. Therefore, 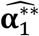 can be computed by back-solving as follows (Strandén and Garrick, 2009; Wang et al., 2012; Bouwman et al., 2017):

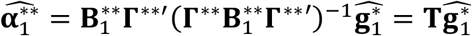

where 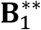 is a diagonal matrix (e.g., an identity matrix **I**) with optional different weights to differentially shrink different loci.

Based on the properties of mixed models (Henderson, 1984), the prediction error covariance matrix of 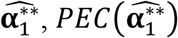, can be obtained as follows:

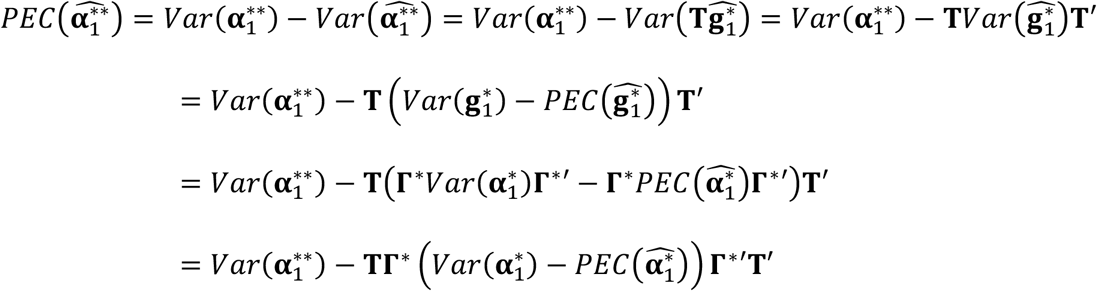

